# SMOC2, OGN, FCN3, and SERPINA3 could be biomarkers for the evaluation of acute decompensated heart failure caused by venous congestion

**DOI:** 10.1101/2024.03.04.583279

**Authors:** Yiding Yu, Huajing Yuan, Quancheng Han, Jingle Shi, Xiujuan Liu, Yitao Xue, Yan Li

**Author notes:** **Correspondence:** Correspondence should be addressed to Yi-tao Xue; and Yan Li;.

## Abstract

**Background:** Venous congestion (VC) sets in weeks before visible clinical decompensation, progressively increasing cardiac strain and leading to acute heart failure (HF) decompensation. Currently, the field lacks a universally acknowledged gold standard and early detection methods for VC.

**Methods:** Using data from the GEO database, we identified VC’s impact on HF through key genes using Limma and STRING databases. The potential mechanisms of HF exacerbation were explored via GO and KEGG enrichment analyses. Diagnostic genes for acute decompensated HF were discovered using LASSO, RF, and SVM-REF machine learning algorithms, complemented by single-gene GSEA analysis. A nomogram tool was developed for the diagnostic model’s evaluation and application, with validation conducted on external datasets.

**Results:** Our findings reveal that VC influences 37 genes impacting HF via 8 genes, primarily affecting oxygen transport, binding, and extracellular matrix stability. Four diagnostic genes for HF’s pre-decompensation phase were identified: SMOC2, OGN, FCN3, and SERPINA3. These genes showed high diagnostic potential, with AUCs for each gene exceeding 0.9 and a genomic AUC of 0.942.

**Conclusions:** Our study identifies four critical diagnostic genes for HF’s pre-decompensated phase using bioinformatics and machine learning, shedding light on the molecular mechanisms through which VC worsens HF. It offers a novel approach for clinical evaluation of acute decompensated HF patient congestion status, presenting fresh insights into its pathogenesis, diagnosis, and treatment.

## INTRODUCTION

Despite adherence to guideline-directed medical therapies, the prognosis for heart failure (HF) patients remains suboptimal. Individuals in advanced stages of HF often necessitate recurrent hospital admissions and enduring pharmacological intervention, imposing significant demands on healthcare resources^1^. The primary catalyst for these hospitalizations is the clinical manifestations of venous congestion (VC). A diminution in venous capacitance coupled with an augmented return of venous blood to the heart escalates preload. This increase in preload, when juxtaposed with compromised cardiac functionality, precipitates a transition from chronic HF to acute decompensated heart failure (ADHF)^2,3^.

HF precipitates fluid retention and VC, transitioning the body from a low-pressure, healthy biological system to a high-pressure, pathological state. This shift forces organs to operate under significantly elevated venous and interstitial pressures, challenging their physiological functions. Traditional views attribute congestion primarily to deteriorating cardiovascular function, with treatment strategies aimed at achieving euvolemic status and congestion alleviation through diuretic therapy^4^. However, emerging evidence suggests that venous congestion may initiate weeks prior to the clinical manifestations of decompensation that necessitate medical intervention^5^. Biomechanical forces, such as increased venous pressure and shear stress, can induce endothelial cells to undergo a phenotypic transformation towards pro-oxidative, pro-inflammatory, and vasoconstrictive states^6^. Biomechanical stress triggers endothelial cells to secrete cytokines, including TNF-α, IL-6, and VCAM-1^7,8^. Yet, the intricate linkage and precise mechanisms underlying endothelial activation due to VC in the context of HF require further investigation. A deeper understanding of these mechanisms is critical for evaluating the clinical efficacy and optimizing the application of diuretic therapies. Concurrently, identifying pivotal biomarkers could pave the way for early detection of VC—a domain where a definitive diagnostic gold standard is currently absent. By identifying and targeting these key markers, there is potential to preemptively shift treatment modalities, thereby reinstating endothelial cell stability and preventing the progression to a decompensated state in heart failure patients.

The swift progress in high-throughput technologies and bioinformatics holds promise for elucidating the mechanisms by which venous congestion triggers endothelial activation in heart failure. Furthermore, the advancement and refinement of machine learning applications within bioinformatics are poised to revolutionize the selection of diagnostic tools. These machine learning algorithms offer the potential to identify sensitive and specific markers, thus furnishing early detection methods for heart failure decompensation.

Based on this, this study analyzed the mRNA data sets of VC and HF in the GEO database and identified differentially expressed genes (DEGs) through the limma package. We performed immune infiltration analysis on the VC dataset to assess whether venous pressure affects immune cell composition. Subsequently, we obtained the interaction relationship between VC and HF differential genes through the STRING database and established a protein-protein interaction network (PPI). Through PPI, we obtained the key genes that HF is affected by VC. We performed enrichment analysis on these genes to obtain the impact of VC on the biological processes and functions of HF. Subsequently, we screened key genes through three machine learning models including least absolute shrinkage and selection operator (LASSO), random forest (RF), and support vector machine recursive feature elimination (SVM-REF). We established a diagnostic model for pre-decompensated heart failure and completed nomogram evaluation and GSEA of individual genes. We also validated the diagnostic model on external datasets. Figure 1 depicts the study flowchart.

**Figure 1:**
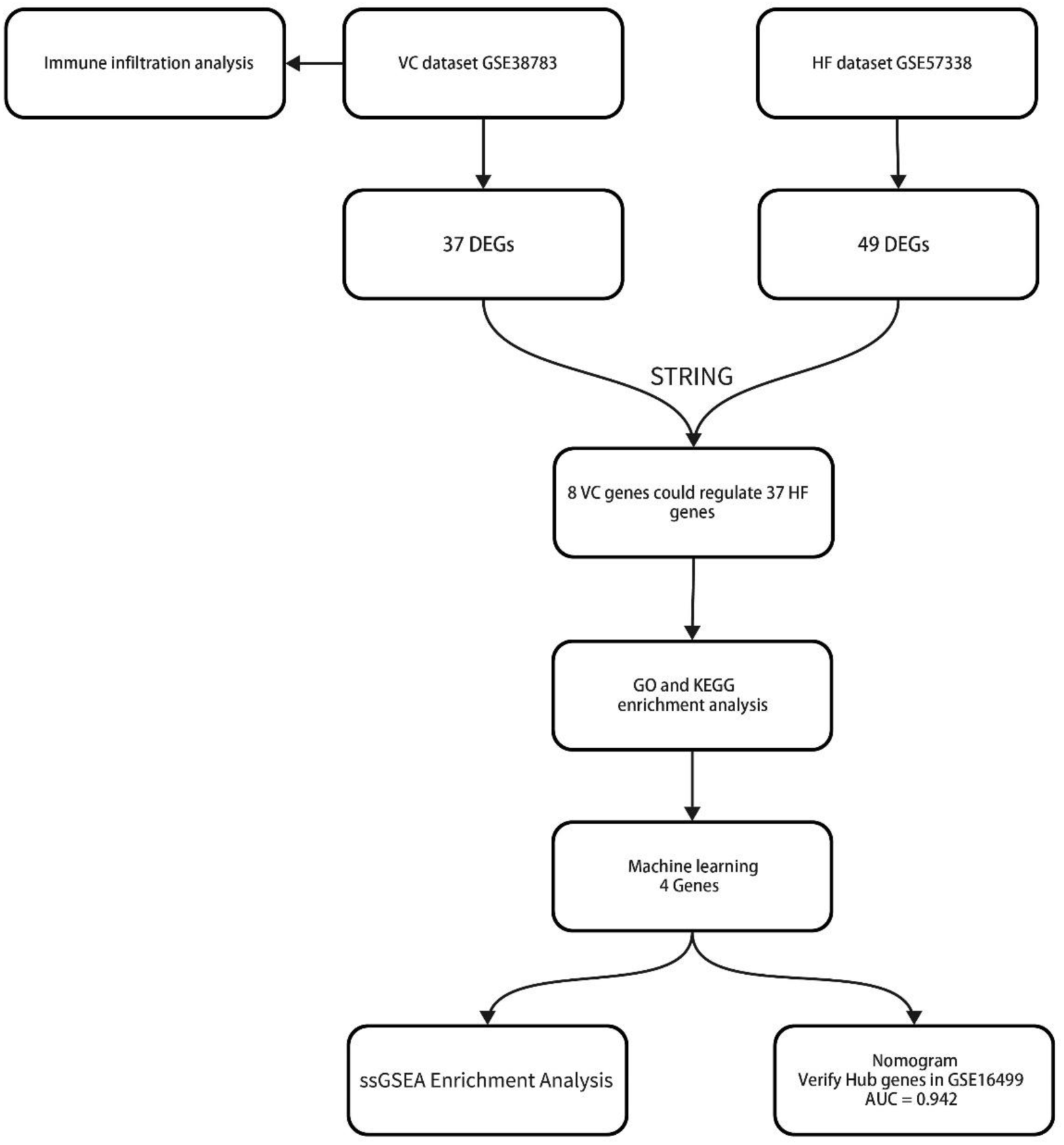
The study flowchart.

## MATERIALS AND METHODS

### Microarray Data

We choose GSE38783 as the VC data set^9^. The trial involved 12 healthy subjects and inflated a pressure cuff around the dominant arm to increase arm venous pressure to approximately 30 mmHg above baseline for 75 minutes. Endothelial cell samples were procured both pre- and post-intervention through an angiocatheter assisted by an intravascular guidewire. For the exploration of heart failure, the GSE57338 dataset was chosen, representing the most comprehensive dataset available in this domain. It comprises left ventricular myocardial specimens from 95 patients diagnosed with ischemic heart failure, alongside 136 control samples from healthy individuals^10^.To corroborate our findings, the GSE16499 dataset served as the validation cohort, encompassing left ventricular myocardial samples from 15 patients with ischemic heart failure and an equivalent number of healthy controls^11^.

### Data Processing and Differentially Expressed Gene Screening

We used R software (R version 4.2.0) to complete data preprocessing. We remove probes corresponding to multiple molecules. For multiple probes corresponding to the same molecule, only the probe with the largest signal value is retained. Finally, we remove the batch effect of the data and convert the probe ID into a gene symbol according to the annotation file of the platform. We used the limma package for differential analysis and selected genes with p-value <0.05 and |log2(FC)|≥1 as differential genes^12^. We use the ggplot2 package to complete the drawing of the picture.

### Immune Infiltration Analysis

Endothelial cell activation will produce cytokines. Therefore, we used the CIBERSORT package to evaluate whether venous pressure has an impact on the endothelial immune environment^13^. Bar charts are used to visualize the proportion of each type of immune cell in different samples. The differences in cell distribution between VC and normal groups were compared by t test, and the cutoff value was set at p<0.05.

### Protein–Protein Interaction Network Construction

In order to understand the process of VC affecting HF and discover the interactions between protein-coding genes, we established a PPI network using the STRING database^14^. Parameter selection default settings. Subsequently, we imported the results into Cytoscape3.6.1 to complete visualization and subsequent analysis^15^.

### Functional Enrichment Analysis

Utilizing the Protein-Protein Interaction (PPI) network, we identified crucial genes implicated in Heart Failure (HF) that are influenced by Venous Congestion (VC). The ’clusterProfiler’ package facilitated the execution of Gene Ontology (GO) and Kyoto Encyclopedia of Genes and Genomes (KEGG) enrichment analyses, enabling the comprehensive visualization and elucidation of these key genes’ roles and pathways^16–18^.

### Machine Learning

To refine the selection of diagnostic markers for the pre-decompensation stage of heart failure, our study employs a triad of machine learning algorithms: Least Absolute Shrinkage and Selection Operator (LASSO), Random Forest (RF), and Support Vector Machine-Recursive Feature Elimination (SVM-RFE)^19–21^. The LASSO algorithm was implemented using the ’glmnet’ package, with ten-fold cross-validation employed to identify the most significant genes. The ’randomForest’ package was utilized to execute the RF algorithm, prioritizing genes based on their importance scores. Similarly, the SVM-RFE algorithm was conducted using the ’e1071’ package, selecting genes that contribute to the highest classification accuracy. After completing the calculation, we selected the intersection of the three as the diagnostic gene for the pre-decompensation stage of heart failure.

### Nomogram Construction and Validation of diagnostic model

For the construction of a diagnostic nomogram for the precompensation phase of heart failure, our analysis leveraged the ’rms’ package, which facilitated the visualization of the contributory weight of each candidate gene as individual points on the nomogram, with ’Total Points’ reflecting the cumulative score derived from all selected genes^22^. Subsequently, boxplots were generated to illustrate the expression profiles of these genes, and Receiver Operating Characteristic (ROC) curves were plotted to assess their diagnostic efficacy. The diagnostic utility was quantified using the Area Under the Curve (AUC), with values exceeding 0.7 deemed indicative of substantial diagnostic merit. To validate our findings, we conducted an analysis on both individual and combined gene sets within the GSE16499 dataset. The discriminatory power of our diagnostic model was further appraised through the ROC curve analysis, ensuring a robust evaluation of its predictive capacity.

### ssGSEA Enrichment Analysis

To elucidate the functional roles of genes identified in the pre-decompensated stage of heart failure, our study employed the ’clusterProfiler’ package to conduct single-gene Gene Set Enrichment Analysis (GSEA)^23^. This analysis was pivotal in uncovering the biological processes and pathways through which these genes may contribute to the transition from chronic to acute decompensated heart failure.

## RESULTS

### Identification of Differentially Expressed Genes

Following a differential analysis, we discerned a total of 37 distinct genes within the VC dataset, comprising 36 up-regulated and a single down-regulated gene. In the HF dataset, we ascertained a total of 49 distinct genes, inclusive of 28 up-regulated and 21 down-regulated genes. Notably, there exists no overlap between the differential genes identified in the two datasets. The illustrative outcomes are depicted in Figure 2.

**Figure 2:**
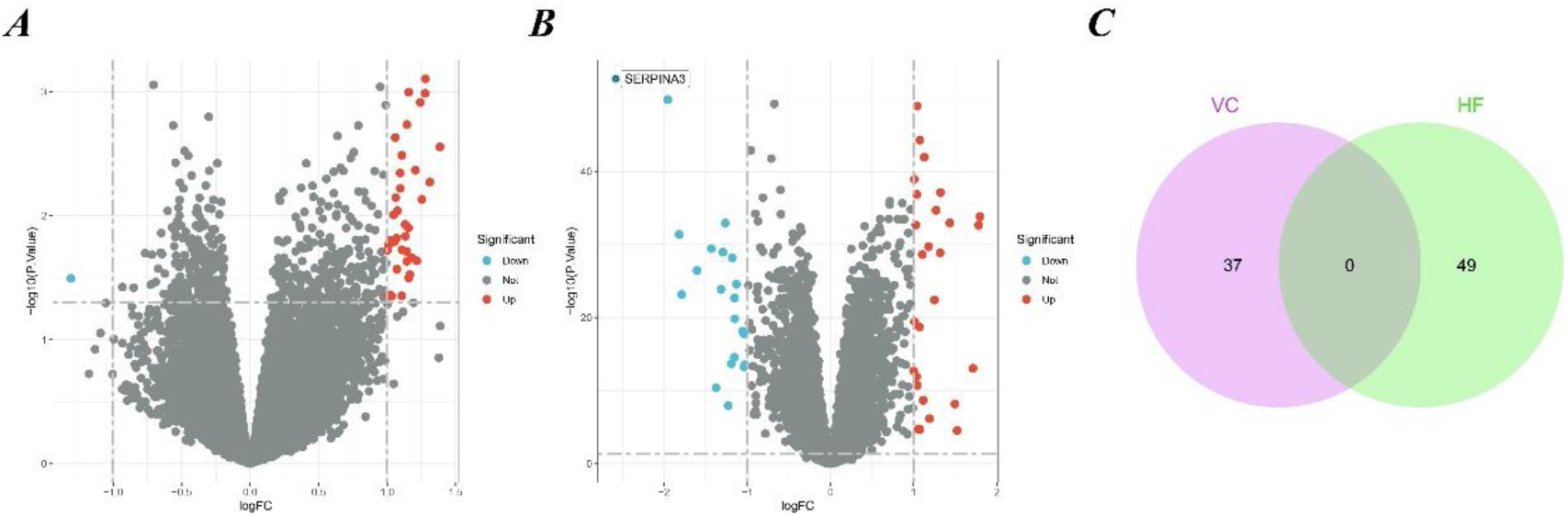
Results of Differentially Expressed Genes. (A) Volcano plot of VC. We set adjust p-values < 0.05 and | log2(FC)| ≥ 1 as the difference genes. (B) Volcano plot of HF. (C) The intersection of VC and HF. We intersected the results to get 0 genes.

### Immune Cell Infiltration Analysis

We performed immune infiltration analysis on the VC data set through the CIBERSORT algorithm. Figure 3 clearly shows the different subpopulation contents within each sample. The results showed that 75 minutes of venous hypertension did not have a significant impact on the immune environment of endothelial cells.

**Figure 3:**
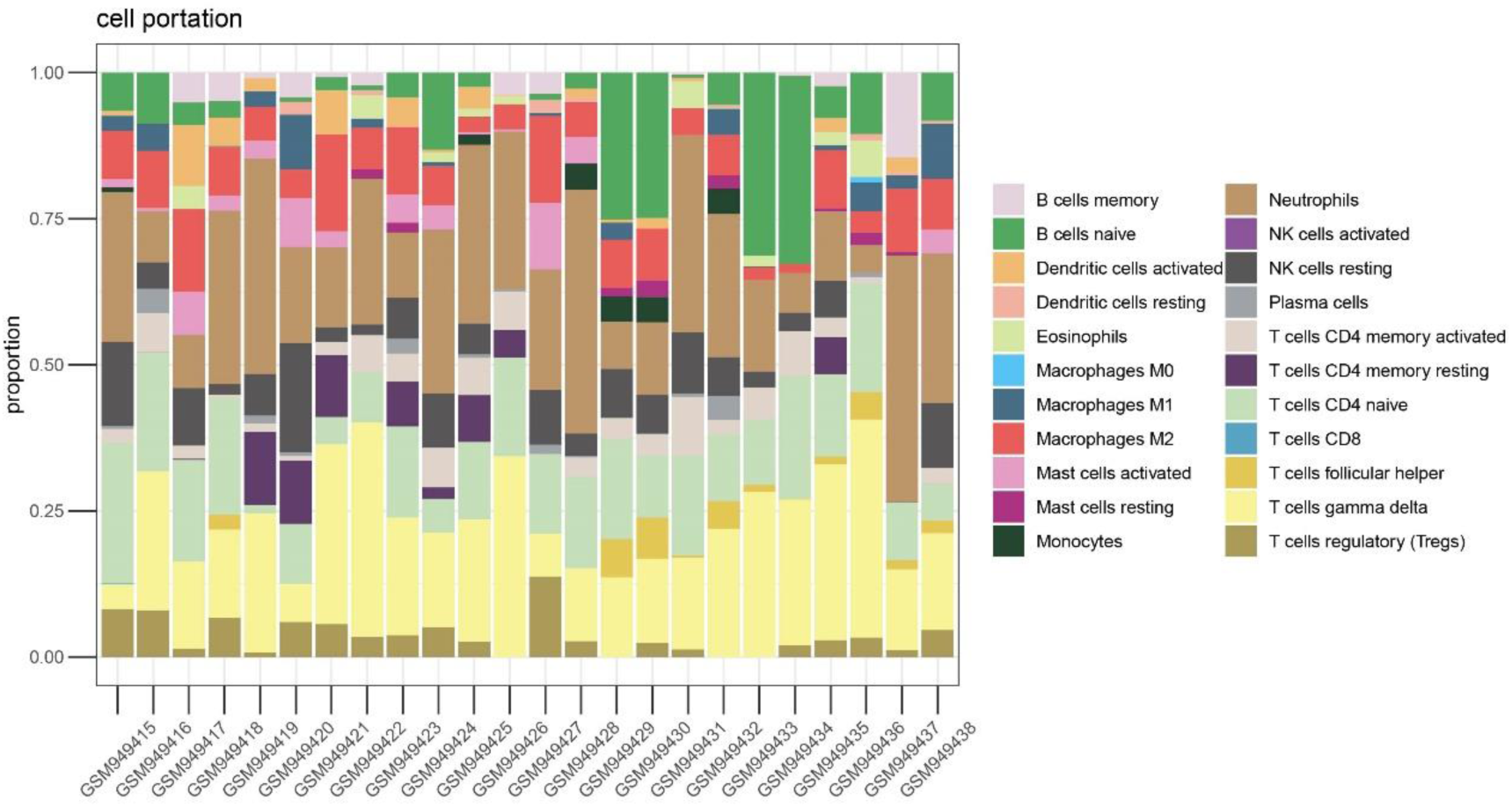
Immune infiltration analysis before and after venous hypertension treatment. The proportion of 22 kinds of immune cells in different samples visualized from the bar plot.

### PPI network construction and hub gene selection

Owing to the lack of overlap between the differential genes of VC and HF, we employed the STRING database to construct a Protein-Protein Interaction (PPI) network, thereby elucidating the interactions among the encoded proteins of VC and HF. Through this PPI network, we pinpointed 45 nodes and 75 interactions. The analysis implied that 8 genes from VC might interact with 37 genes from HF. The visualization results are shown in Figure 4.

**Figure 4:**
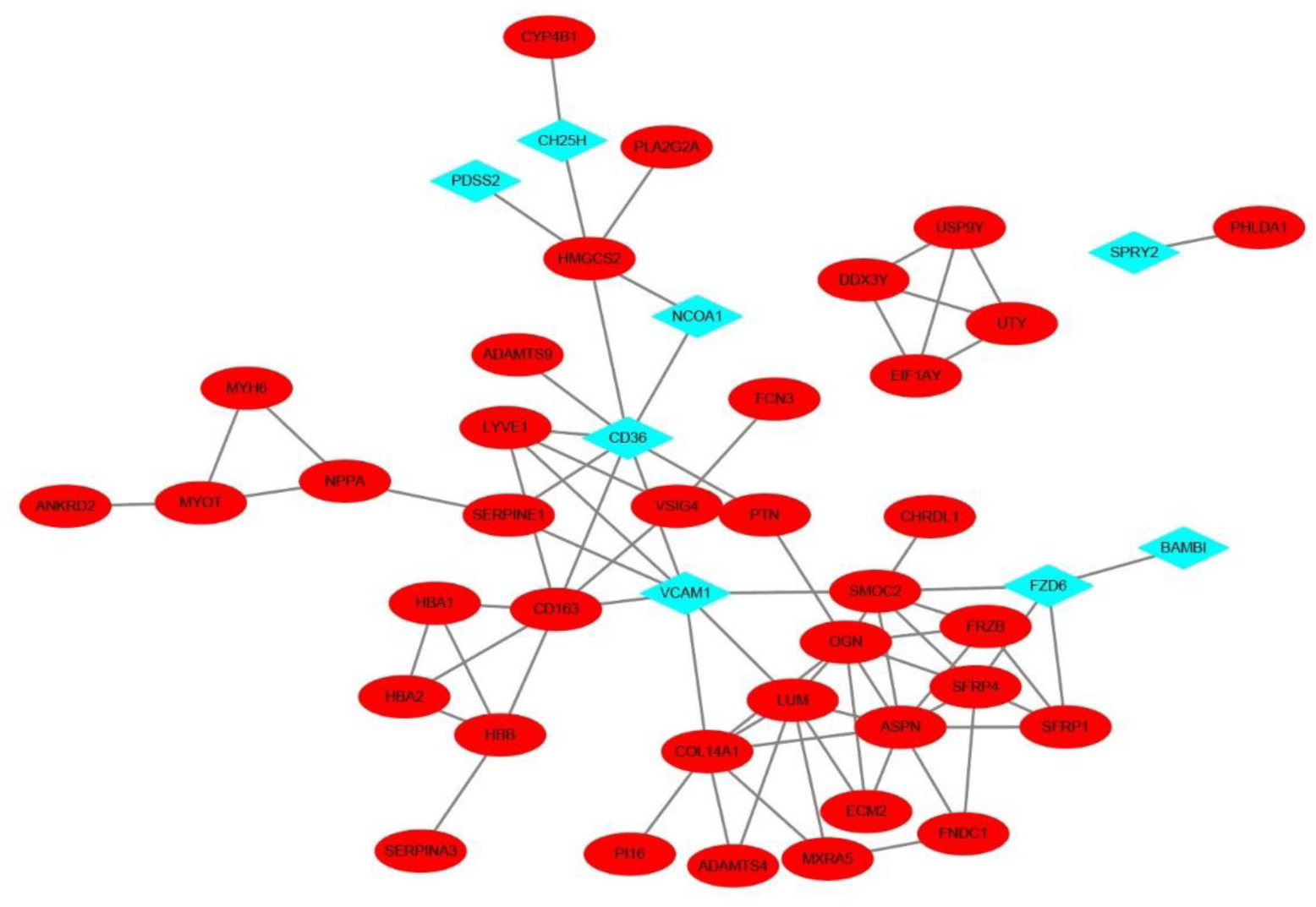
PPI network of VC and HF. The blue diamond-shaped node represents the VC gene, and the red oval-shaped node represents the HF gene.

### GO and KEGG enrichment analysis

To evaluate the impact of VC on HF biological processes and functions, we performed GO and KEGG enrichment analysis on 37 HF genes. Categories of GO analysis include biological processes (BP), cellular components (CC), and molecular functions (MF). The predominantly enriched BP terms encompassed those related to oxygen transport, gas transport, extracellular matrix organization, extracellular structure organization, and the organization of external encapsulating structures. The chiefly enriched CC terms included the collagen-containing extracellular matrix, the haptoglobin-hemoglobin complex, the hemoglobin complex, blood microparticles, and the lumen of endocytic vesicles. The MF terms that were most enriched involved haptoglobin binding, oxygen binding, oxygen carrier activity, collagen binding, and the extracellular matrix structural constituent conferring compression resistance. In the KEGG pathway enrichment analysis, these 37 genes were primarily enriched in pathways associated with African trypanosomiasis, Malaria, the Wnt signaling pathway, and the Complement and coagulation cascades. The enrichment results were shown in Figure 5.

**Figure 5:**
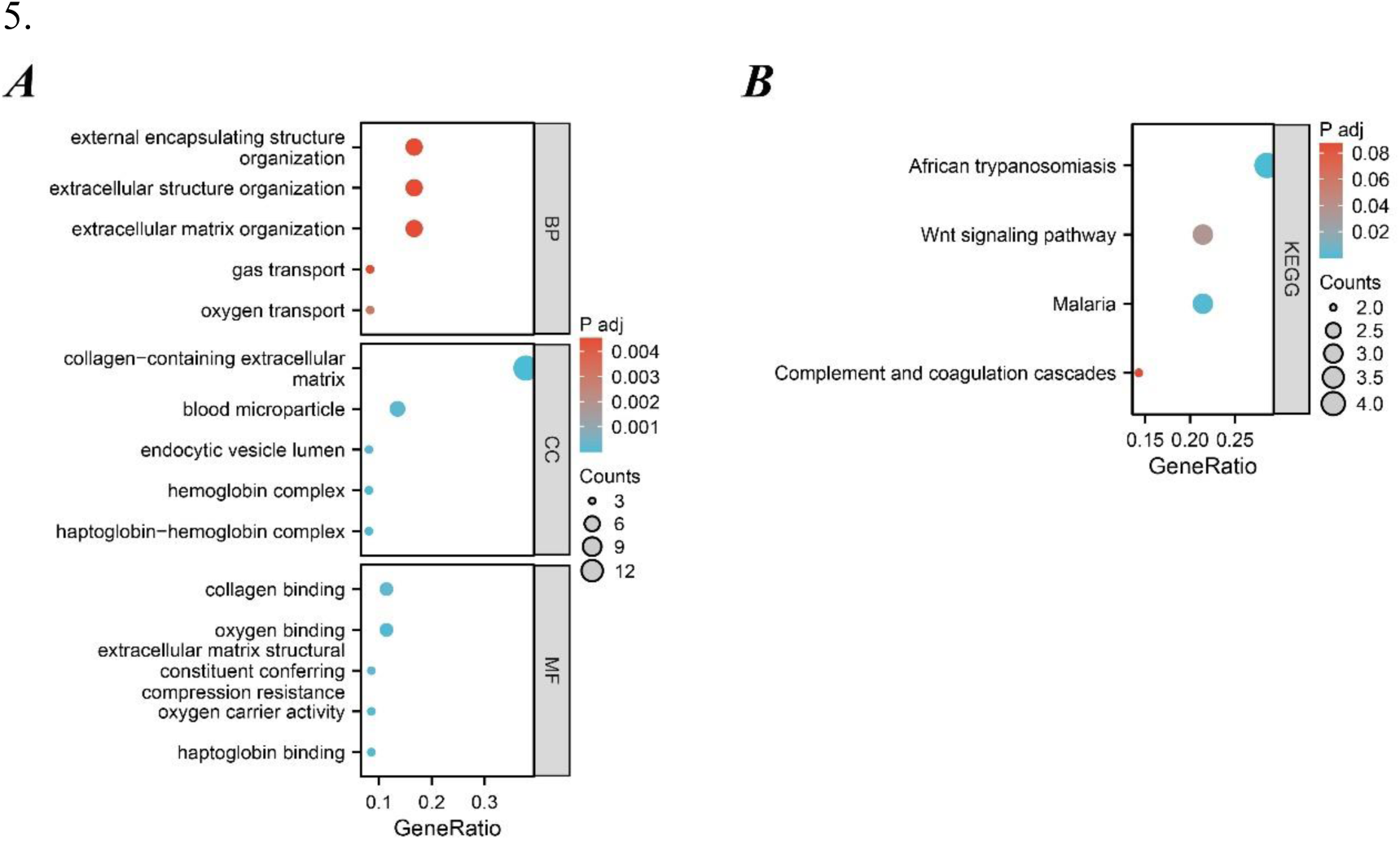
Functional Enrichment Analysis of 37 HF genes. (A) GO enrichment analysis results. (B) KEGG enrichment analysis results.

### Identification of Hub Genes via Machine Learning

We employed three machine learning algorithms—LASSO, RF, and SVM-RFE—to identify diagnostic genes for the pre-decompensation stage of heart failure. The LASSO algorithm revealed 15 potential genes, while the RF algorithm prioritized genes based on their calculated significance, selecting the top six genes with the utmost importance as candidates. The SVM-RFE algorithm indicated optimal accuracy with 29 genes; hence, we adopted the initial 29 genes from the SVM-RFE output as candidate genes. By intersecting the findings from all three algorithms, we pinpointed four diagnostic genes for the pre-decompensation stage of heart failure: SMOC2, OGN, FCN3, and SERPINA3. These results are displayed in Figure 6.

**Figure 6:**
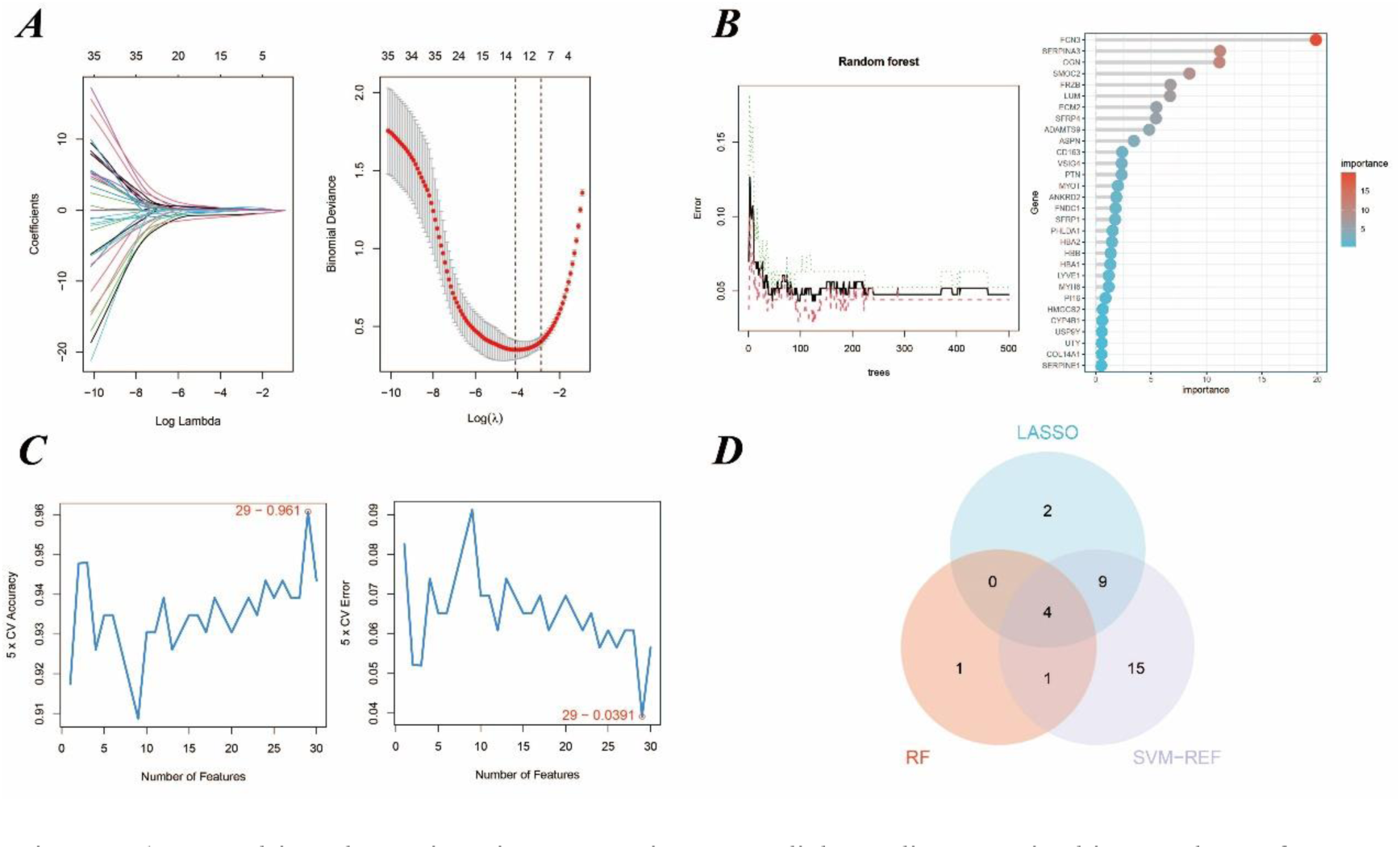
Machine learning in screening candidate diagnostic biomarkers for pre-decompensation stage of HF. (A) In the Lasso model for biomarker screening, we plotted LASSO coefficient profiles of the selected genes, determining the optimal lambda at the point where the model’s error was minimal. Each line in the left graph represents a gene, while the right graph’s vertical lines indicate the model’s error, pinpointing the best gene selection count at 15.(B) The SVM-RFE model’s screening involved utilizing the algorithm to identify the most accurate genes, demonstrated by the plot where the number of genes selected is balanced against their predictive accuracy.(C) In the RF model, we focused on the relative importance of the candidate genes, specifically highlighting the top six genes as determined by the random forest’s importance calculation.(D) A Venn diagram consolidates the outcomes, indicating four genes consistently identified across the three analytical methods.

### Diagnostic Value Assessment

We devised a nomogram anchored on four pivotal hub genes, accompanied by the formulation of a Receiver Operating Characteristic (ROC) curve, to meticulously gauge the diagnostic specificity and sensitivity conferred by each gene and the nomogram as a whole. Furthermore, we illustrated the expression disparities of these hub genes within the heart failure (HF) dataset via a box plot. Subsequent validation of the hub gene on the external dataset GSE16499 demonstrated robust diagnostic efficacy, with the Area Under the Curve (AUC) for each gene surpassing 0.9 and the collective genomic AUC reaching 0.942, underscoring significant diagnostic potential. The visual representation of these findings was encapsulated in Figure 7.

**Figure 7:**
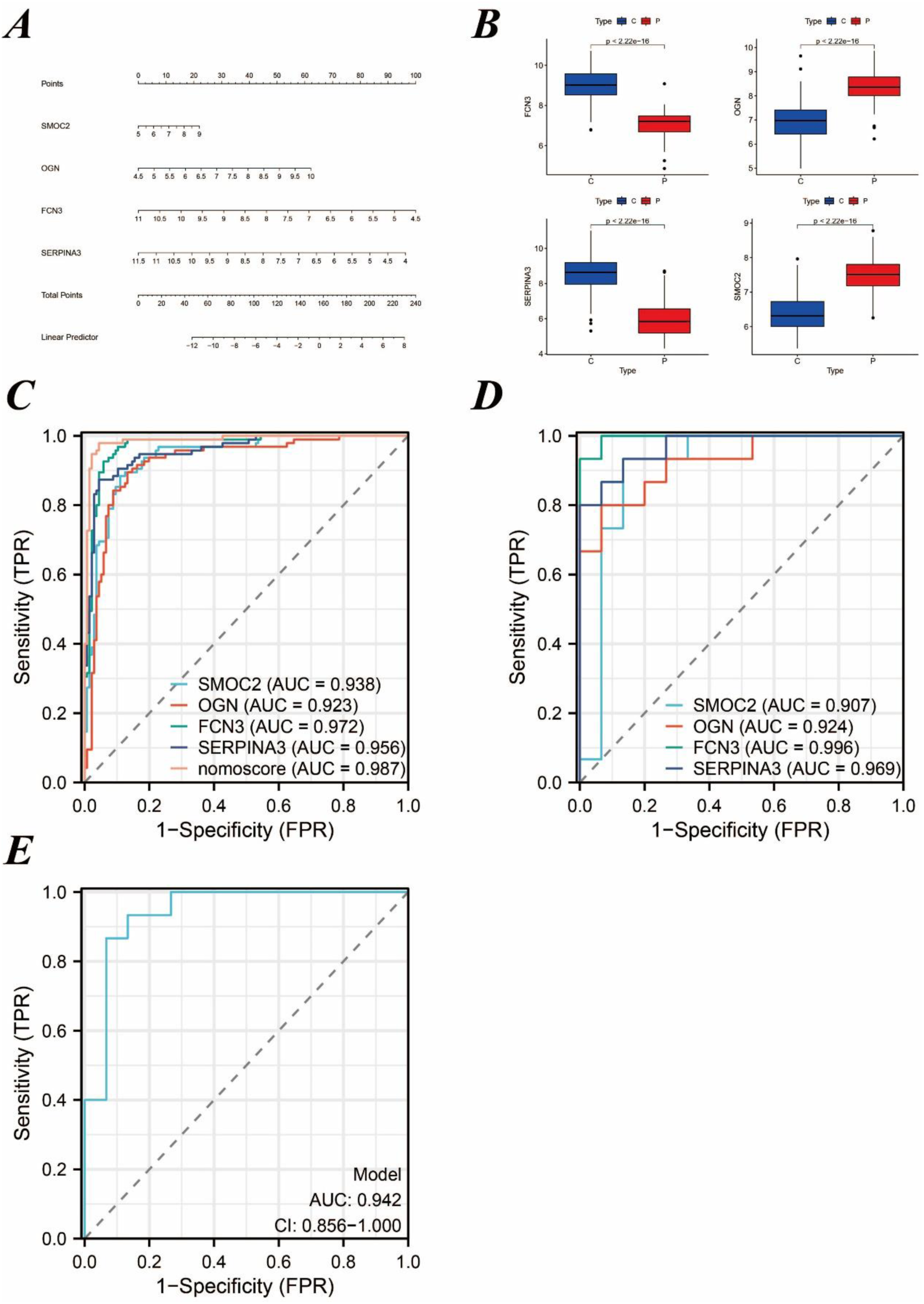
Results of Diagnostic Value Assessment. (A) The illustrative nomogram for diagnosing pre-decompensation stage of HF.(B) Differential expression of hub genes in HF patients relative to normal controls in dataset GSE57338. (C) The ROC curve for the nomogram and individual candidate genes, highlighting their diagnostic value for pre-decompensation stage of HF. (D) The ROC curves for each candidate gene within the GSE16499 dataset. (E) The ROC curve representing the diagnostic efficacy of the 4-gene model in dataset GSE16499.

### Single-sample Gene Set Enrichment Analysis

We performed ssGSEA on SMOC2, OGN, FCN3, and SERPINA3, respectively. The analysis revealed that these genes contribute variably to the biosynthesis and metabolism of amino acids, the production of steroid compounds, and the synthesis of nucleotide sugars. Comprehensive results and their visual representations are detailed in Figure 8.

**Figure 8:**
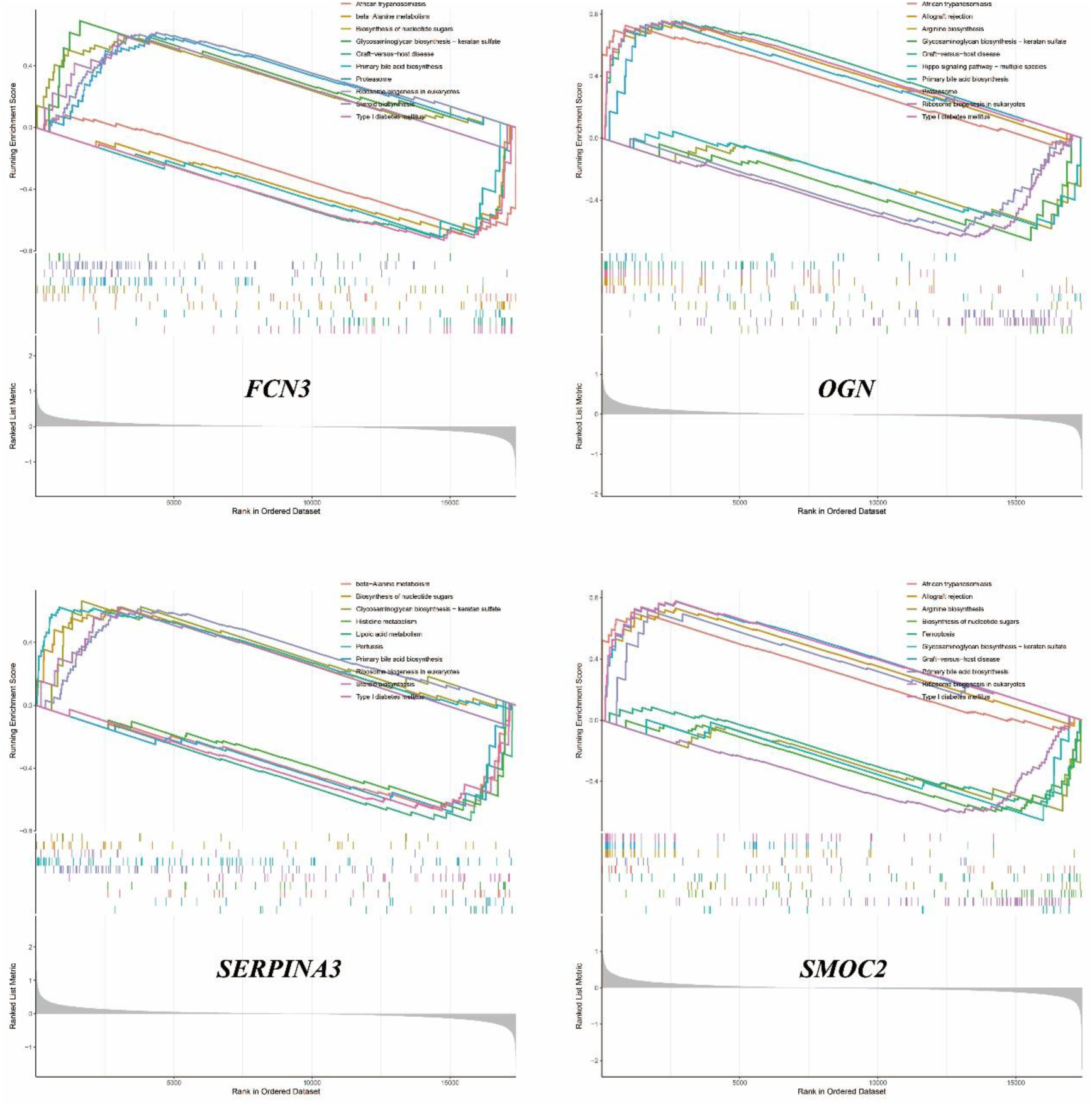
The results of ssGSEA.

## DISCUSSION

Venous congestion induces endothelial and neurohormonal activation, signifying a decline in cardiac function. This activation is not only a consequence but also a catalyst for further hemodynamic decline in heart failure, often precipitating the transition from chronic to acute decompensated heart failure. Consequently, venous congestion emerges as a crucial hemodynamic indicator for predicting rehospitalization and mortality post-discharge in patients with acute decompensated heart failure^24^. However, it’s pivotal to recognize that venous congestion manifests in the initial phase of decompensation. Deciphering its impact on cardiac function and identifying early diagnostic markers are essential for timely modulation of therapeutic strategies.

In this study, we first obtained the differential genes of VC and HF through differential analysis. Regrettably, there was no overlap between the differential genes for VC and HF, suggesting that the pathogenesis of heart failure during stable periods devoid of congestion might not be directly linked to VC. However, utilizing the Protein-Protein Interaction (PPI) network, we discovered that subsequent to the onset of VC, eight differential genes in the vascular endothelium could potentially influence 37 genes associated with HF. These 37 genes might be pivotal in the transition of heart failure patients from a stable condition to the acute decompensated stage.

We explored the biological functions and signaling pathways involved in key genes through GO and KEGG enrichment analysis. The results show that the mechanism by which VC causes acute decompensation of heart failure may be related to oxygen transport and oxygen binding. Heart failure is characterized by exercise intolerance, which is closely related to structural and functional abnormalities in oxygen transport^25^. When heart function decreases, the heart cannot pump blood effectively and meet the body’s demand for oxygen^26^. It can be seen from the PPI network that after the occurrence of VC, VCAM1, CD36 and NCOA1 of endothelial cells change, affecting the HAB1, HAB2, and HABB genes of HF, which may further reduce oxygen transport and oxygen binding and other functions. Hemoglobin plays a key physiological role in transporting oxygen in the human body. However, VC may increase levels of haptoglobin, an acute-phase serum protein that binds very strongly and irreversibly to hemoglobin and is endocytosed by macrophages^27^. At the same time, CD36 and VCAM1 may induce an increase in CD163 levels after endothelial activation, degrade the hemoglobin-haptoglobin complex, and produce an inflammatory response^28^.The enrichment analyses further suggest that VC may influence collagen binding and the organization of the myocardial extracellular matrix. This matrix serves as a sophisticated, evolving scaffold that upholds tissue structure and function^29^. Remodeling of the extracellular matrix can hinder cardiac filling, exacerbating heart failure through an inflammatory response^30^. Moreover, this remodeling process can alter the collagen content, impacting the heart chamber’s stiffness. Variations in the quantity and quality of collagen can significantly affect myocardial stiffness. Persistent elevation in wall stress, exceeding the heart’s adaptive capacity, may result in a disproportionate reduction in ventricular wall thickness relative to the ventricular volume, culminating in diastolic and systolic dysfunction^31^.

The emergence of venous congestion typically precedes the clinical signs of decompensation by weeks, often culminating in overt clinical deterioration that necessitates medical intervention. To address this, we employed a machine learning algorithm to identify four diagnostic genes associated with ADHF triggered by VC, namely SMOC2, OGN, FCN3, and SERPINA3. A nomogram was developed to enhance the practical utility of the diagnostic model. These four genes were subsequently validated against the external dataset GSE16499, where the AUC for each gene exceeded 0.9, and the composite genomic AUC was 0.942, underscoring their significant diagnostic potential. We completed the ssGSEA of these four genes, hoping to provide new ideas for our understanding of the molecular mechanism of ADHF after VC.

SPARC-related modular calcium-binding protein 2 (SMOC2), part of the SPARC family of matricellular proteins, is known to promote endothelial cell proliferation, migration, and angiogenesis^32^. Research indicates that SMOC2 expression escalates in rats experiencing heart failure, and silencing SMOC2 can mitigate heart failure symptoms by modulating autophagy via the TGF-β1/Smad3 signaling pathway^33^. Additionally, mouse studies have demonstrated that SMOC2 knockdown can ameliorate cardiac function impairment and cardiac fibrosis, potentially through the inhibition of the ILK/p38 signaling pathway^34^. Furthermore, the overexpression of SMOC2 has been observed to enhance vascular smooth muscle cell proliferation, migration, and extracellular matrix breakdown, pointing to its significant role in cardiovascular pathophysiology^35^.

Osteoglycin (OGN), a constituent of the small leucine-rich repeat proteoglycan (SLRP) family, plays a critical role in modulating inflammation and cardiac fibrosis^36^. Research has revealed that OGN attenuation can suppress cardiac fibroblast proliferation and the epithelial/endothelial-mesenchymal transition, pivotal processes in cardiac remodeling. Moreover, OGN is known to facilitate apoptosis in cardiac fibroblasts, mediated through the regulation of the Wnt signaling pathway^37^. Variations in OGN levels offer distinct insights into myocardial remodeling dynamics^38^. Notably, an upsurge in OGN expression within infarct scars contributes to the proper maturation of collagen, thereby preventing cardiac rupture and counteracting adverse remodeling post-myocardial infarction (MI). Consequently, OGN holds promise as a valuable biomarker for ischemic heart failure^39^.

Ficolins constitute a family of proteins characterized by their collagen-like and fibrinogen-like domains, playing a crucial role in triggering the complement lectin pathway^40^. Ficolin-3 (FCN3), a principal molecule in this pathway, is pivotal for the activation of complement component 3 and has been linked to hypertension risk^41^. Emerging evidence from studies on patients with acute congestive heart failure suggests an association with complement activation, positioning FCN3 could as a potential biomarker for this condition^42^. However, the results of two clinical studies indicate that complement activation products have no prognostic value in acute heart failure^43,44^. Furthermore, the activation of the complement system and the consequent production of reactive oxygen species in the myocardium post-infarction are known to stimulate the upregulation of cytokines and chemokines. This cascade can precipitate adverse outcomes, such as left ventricular dilatation and contractility impairments^45^.

Serpin Family A Member 3 (SERPINA3), also known as α-1-antichymotrypsin (AACT or ACT), is part of the serpin superfamily, which primarily functions to inhibit serine proteases. This protein’s elevated levels are observed in conditions such as heart failure and various neurological disorders, including Alzheimer’s disease and Creutzfeldt-Jakob disease^46^. Research indicates that SERPINA3 acts as an endogenous gene responsive to the mineralocorticoid receptor and can be upregulated by aldosterone^47^. Following VC, alterations in fluid retention and mineralocorticoid levels might disrupt SERPINA3 balance. Clinical evidence suggests that heightened SERPINA3 levels in individuals with new-onset or exacerbating heart failure correlate with increased mortality rates or unplanned cardiac readmissions, underscoring its prognostic value for heart failure^48^. Furthermore, SERPINA3 has been proposed as a biomarker for assessing right ventricular myocardial function, particularly in the context of systemic congestion leading to right ventricular failure^49^. Beyond its myocardial expression, elevated circulating levels of SERPINA3 have been linked to an increased cancer risk in heart failure patients, highlighting the necessity for more in-depth exploration of its clinical implications and molecular functions^50^.

There is currently no gold standard for detecting congestion in established acute decompensated heart failure. This is usually assessed by signs and symptoms of heart failure, as well as objective evidence of congestion (eg, elevated BNP, pulmonary edema on chest x-ray, or pulmonary rales on auscultation). There is also no uniform or readily available method for determining when effective dehydration occurs or when a patient has reached optimal volume status^51^. This is also the question our research attempts to address.

The novel contributions of our investigation are manifold. Primarily, this study pioneers the bioinformatics exploration of the interplay between venous congestion and heart failure, dissecting the intricate mechanisms through which venous congestion may precipitate the transition from chronic to acute decompensated heart failure. Secondly, we have pinpointed four diagnostic genes—SMOC2, OGN, FCN3, and SERPINA3— associated with acute decompensated heart failure, employing three distinct machine learning techniques for their identification. Validation of these genes confirmed that their collective diagnostic model possesses substantial value, offering a new avenue to assess patient congestion status effectively. Furthermore, our analysis extends into the biological processes influenced by venous congestion in heart failure, providing fresh insights into its potential mechanisms and refining our understanding of the therapeutic implications of diuretics.

Despite its contributions, our study acknowledges certain limitations. Initially, the causal link between the observed elevation in mRNA levels and a corresponding increase in protein expression remains undetermined. Additionally, the possibility that endothelial activation post-venous congestion might trigger the secretion of other molecules influencing heart failure, such as exosomes, has not been fully explored.

While we have developed a nomogram to facilitate the implementation of the diagnostic model, the precise values and their clinical relevance require validation through subsequent clinical trials.

## CONCLUSION

We performed bioinformatics analysis of the GEO dataset to explore the potential molecular mechanisms by which venous congestion affects heart failure. Through three machine learning algorithms, LASSO, RF and SVM-RFE, we identified SMOC2, OGN, FCN3, and SERPINA3 as potential biomarkers for the evaluation of acute decompensated heart failure caused by venous congestion. More importantly, diagnostic models and nomogram tools based on these 4 genes were developed to help clinically assess the hyperemic status of patients with acute decompensated heart failure, which may become an interesting target for future in-depth research.

## Acknowledgements

Not applicable.

## CRediT authorship contribution statement

Yiding Yu: Conceptualization, Methodology, Validation, Formal analysis,

Investigation, Resources, Visualization, Writing – original draft, Writing – review & editing.

Huajing Yuan: Methodology, Validation, Writing – review & editing. Quancheng Han: Investigation, Resources, Writing – review & editing. Jingle Shi: Supervision, Software, Writing – review & editing.

Xiujuan Liu: Supervision, Software, Data curation, Writing – review & editing.

Yitao Xue: Project administration, Writing – review & editing.

Yan Li: Project administration, Writing – review & editing, Funding acquisition.

## Consent for publication

Not applicable.

## Ethics approval and consent to participate

Not applicable.

## Data availability statement

Publicly available datasets were analyzed in this study. This data can be found here: GSE38783; GSE57338; GSE16499.

